# Heat-stable preservation of protein expression systems for portable therapeutics production

**DOI:** 10.1101/089664

**Authors:** David Karig, Seneca Bessling, Peter Thielen, Sherry Zhang, Joshua Wolfe

**Affiliations:** Research and Exploratory Development Department, Johns Hopkins University Applied Physics Laboratory, Laurel, MD, USA; Department of Chemical and Biomolecular Engineering, Johns Hopkins University, Baltimore, MD, USA

## Abstract

Many biotechnology capabilities are limited by stringent storage needs of reagents, largely prohibiting use outside of specialized laboratories. Focusing on a large class of protein-based biotechnology applications, we address this issue by developing a method for preserving cell-free protein expression systems under months of heat stress. Our approach realizes an unprecedented degree of long term heat stability by leveraging the sugar alcohol trehalose, a simple, low-cost, open-air drying step, and strategic separation of sets of reaction components during drying. The resulting preservation capacity opens the door for efficient production of a wide range of on-demand proteins under adverse conditions, for instance during emergency outbreaks or in remote or otherwise inaccessible locations. As such, our preservation method stands to advance a great number of different cell-free technologies, including remediation efforts, point of care therapeutics, and large-scale biosensing. To demonstrate this application potential, we use cell-free reagents subjected to months of heat stress and atmospheric conditions to produce sufficient concentrations of a pyocin protein to kill *Pseudomonas aeruginosa*, one of the most troublesome pathogens for traumatic and burn wound injuries. Our work makes possible new biotechnology applications that demand both ruggedness and scalability.

## INTRODUCTION

The vast majority of biotechnology and synthetic biology capabilities remain limited to the laboratory due to issues with reagent stability, robustness, and safety concerns. This is particularly true of protein-based applications, which often capitalize on the inherent properties of natural products, but remain challenging from a storage perspective. Recently, cell-free protein expression systems have been suggested as a promising path for protein-based applications, as they simplify protein expression, purification, and screening while offering safety advantages over engineered living cells^1-8^. However, cell-free reaction components still suffer from preservation concerns, typically requiring cold-chain storage. The ability to preserve and store protein production machinery, particularly at and above room temperature, would drive a breadth of new applications, enabling therapeutics, biosensors, and remediation approaches^9, 10^, including in emergency situations or remote field locations where delivery of lab synthesized products is infeasible.

To pave a path towards these applications, we sought to develop an approach that meets all of the following criteria: 1) preservation of all components needed for transcription and translation, 2) long-term heat stability, 3) potential for scalability, 4) stability under atmospheric conditions, and 5) ability to produce an active therapeutic using stored reagents. The preservation of protein expression systems is challenging because these systems consist of around one hundred essential proteins and small molecules.^11^ Previously, several important efforts have taken steps towards realizing some, but not all of the above criteria. In particular, there is a lack of methods to date that demonstrate long term heat stability.

Two early cell-free preservation efforts focused on drying wheat germ translation systems with stabilizers. These studies, however, did not demonstrate long-term heat stabilization (e.g. months at 37°C).^12, 13^ More recently, Pardee et al. took a major step towards fieldable protein expression systems when they lyophilized cell-free reagents and used them to both implement biosensor gene networks capable of detecting Ebola^14^ and Zika virus^15^ and also to produce a wide array of therapeutics^16^. Although the Pardee et al. methods demonstrate stabilization at room temperature, they still do not fully address the ruggedness needs for many applications. First, long-term (one year) stability for their pellet-based approach was demonstrated using an inert gas atmosphere (N_2_) to prevent oxidative damage, and a silicon desiccant package to prevent hydrolytic damage^14^. Second, their paper-based approach for short-term (24 h) storage is excellent for many small-scale biosensing applications, but not for larger scale applications (e.g., remediation or therapeutics) requiring high volumes of extract. Finally, significant heat resilience above room temperature was not demonstrated^13, 17^. Another important work by Smith et al similarly presented the testing of preserved cell extract stored for different durations at room temperature. In this study, however, either fresh energy system reagents (e.g. phosphoenolpyruvate, ammonium glutamate, potassium glutamate, potassium oxalate, NAD, CoA, nucleotide triphosphates, folinic acid, and tRNA) were added at each timepoint, or the stability of dried energy system reagents at room temperature was characterized using fresh extract^17^. Given that both the extract and energy system exhibit pronounced declines in efficacy over two months, it is unclear how a full set of preserved essential cell-free system components would perform. In addition, in this approach, amino acids, spermidine, and putrescine were added separately, and the stability of these reagents was not presented. Finally, reconstitution of the energy system in this design necessitated the addition of optimal amounts of sodium hydroxide, requiring testing with litmus paper. An interesting extension of the Smith et al. approach was recently presented as well. Lyophilized extract was tested at 25°C over the course of a year. Again, however, the addition of fresh energy system reagents appeared to be required.^18^ Moreover, dried, but not shelf-stored extract was used to produce a 12 kDa therapeutic protein, onconase.^18^ Production of onconase on demand would require addition of tRNA at 15-30 minute intervals, and the stability of tRNA was not discussed. Finally, as with the Pardee et al study, stabilization above room temperature was not demonstrated. Thus, while each of the above studies offers important benefits and tradeoffs, a key missing element from all of them is the demonstration of heat stabilization.

Here, we address the limitations of previous preservation systems by introducing a method for storage of complete protein expression kits over months of heat stress under otherwise unregulated atmospheric conditions (Fig. 1a). We then demonstrate the application potential of our approach by using preserved, heat-stressed reagents to produce sufficient concentrations of a pyocin protein to kill the opportunistic human pathogen *Pseudomonas aeruginosa*.

**Figure 1:**
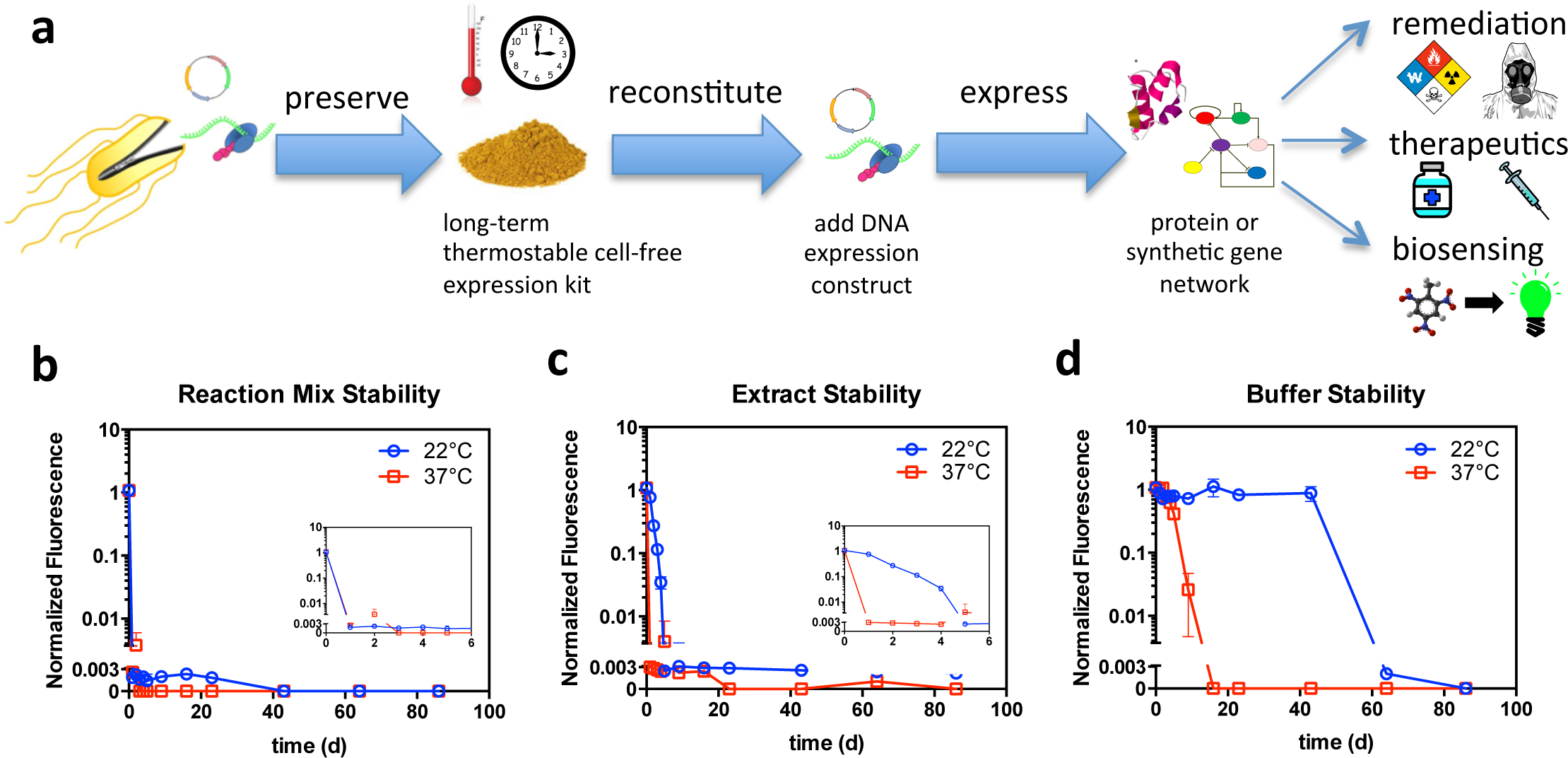
Cell-free preservation overview and baseline reagent stability characterization. (**a**) Several applications stand to benefit from the ability to preserve cell-free protein expression reagents in a scalable, heat and atmosphere stable fashion. Following reconstitution of preserved cell-free reagents, DNA encoding the expression of a protein or gene network of interest is added to enable applications in bioremediation, on demand therapeutics, and large scale biosensing. (**b**) Stability characterization of cell-free reaction mixture. Aliquots of *E. coli* based cell-free reaction mixtures were stored at 22°C and at 37°C. At different time points, expression capacity of the stored mixtures was assessed by adding a T7-EGFP expression construct and measuring fluorescence after 5 h of incubation. The inset zooms in on the first 6 d. (**c**) Stability characterization of *E. coli* cell extracts. Aliquots of extract were stored at 22°C and at 37°C. At different time points, fresh reaction buffer was added, and expression capacity was assessed by adding a T7-EGFP expression construct and measuring fluorescence after 5 h of incubation. The inset zooms in on the first 6 d. (**d**) Stability characterization of reaction buffer. Aliquots of reaction buffer were stored at 22°C and at 37°C. At different time points, fresh extract was added, and expression capacity was assessed by adding a T7-EGFP expression construct and measuring fluorescence after 5 h of incubation. In (b)-(d), we show the means of triplicate fluorescence measurements after 5 h of incubation, with error bars indicating standard deviation. Cases where no error bars are visible indicate that error bars are smaller than the marker. Fluorescence values are normalized by dividing by the 5 h fluorescence of a standard reaction with fresh reagents. The axis break indicates the threshold for significance above background levels.

## RESULTS

We first produced cell-free reagents and defined an assay for quantifying expression capacity. Specifically, we produced *E. coli* extracts using sonication^19^ and leveraged a simplified cell-free reagent production protocol (see Methods)^20^. To quantify expression, we measured fluorescence resulting from expression of enhanced green fluorescent protein (EGFP) from the pUC-T7tet-T7term plasmid. Although we normalize all subsequent fluorescence measurements to a standard experiment with fresh reagents (see Methods), we first compared the efficacy of our reagents to a commercial kit in order to provide a frame of reference (Supplementary Fig. 1). This analysis showed that our reagents outperformed a comparable commercial system by 60%. Having produced reagents and defined an expression assay, we then characterized the baseline stability of our cell-free reaction mixture through daily monitoring of the mixtures’ ability to express enhanced green fluorescent protein (EGFP) from the pUC-T7tet-T7term plasmid. After only a single day at either room temperature or 37°C, unpreserved reaction mixture failed to generate fluorescence (Fig. 1b). Next, we used the same approach to separately characterize the stability of the two main components of the reaction mixture, namely the cell extract and the reaction buffer. For this, days old cell extract was combined with fresh reaction buffer, while days old reaction buffer was combined with fresh cell extract. The EGFP expression plasmid was then added to each sample and the samples were assayed. After 4 d at room temperature, cell extract efficacy had decreased by 31.1-fold. After only 1 d under 37°C heat stress, the extract had completely lost viability and generated negligible fluorescence in our expression assay (Fig. 1c). By contrast, at room temperature, buffer efficacy remained high for 43 d and then rapidly decreased (Fig. 1d). However, after only 9 d at 37°C, buffer efficacy was reduced by 41.5-fold.

Next, we tested the stability of open-air dried reagents. For this, we stored magnesium acetate (a known deliquescent) separately and only combined it with other reagents at the time of assay. Magnesium acetate was separated to enhance stability of dried reagents to hydrolytic damage^21^. Stability quantification experiments were conducted similarly to the initial baseline characterizations. Specifically, (i) dried reaction mixture was reconstituted with water to the appropriate volume, (ii) dried cell extract was reconstituted and combined with fresh reaction buffer and (iii) dried reaction buffer was reconstituted and combined with fresh cell extract. Finally, the EGFP expression plasmid and magnesium acetate were added to each sample and the samples were assayed. Interestingly, while dried reaction mixture completely lost viability after 1 d (Fig. 2a), individually stored extract and reaction buffer still exhibited significant protein expression capacity even after 3 d (Fig. 2b-c). This suggested a potential benefit to separately preserving extract and reaction buffer. Also, although extract was less stable than reaction buffer in liquid form (Fig. 1c-d), dried extract was more stable than dried reaction buffer (Fig. 2b-c). Specifically, even after 40 d, dried extract exhibited less than a 10-fold decrease in expression yield, whereas dried reaction buffer completely lost viability after 6 d.

**Figure 2:**
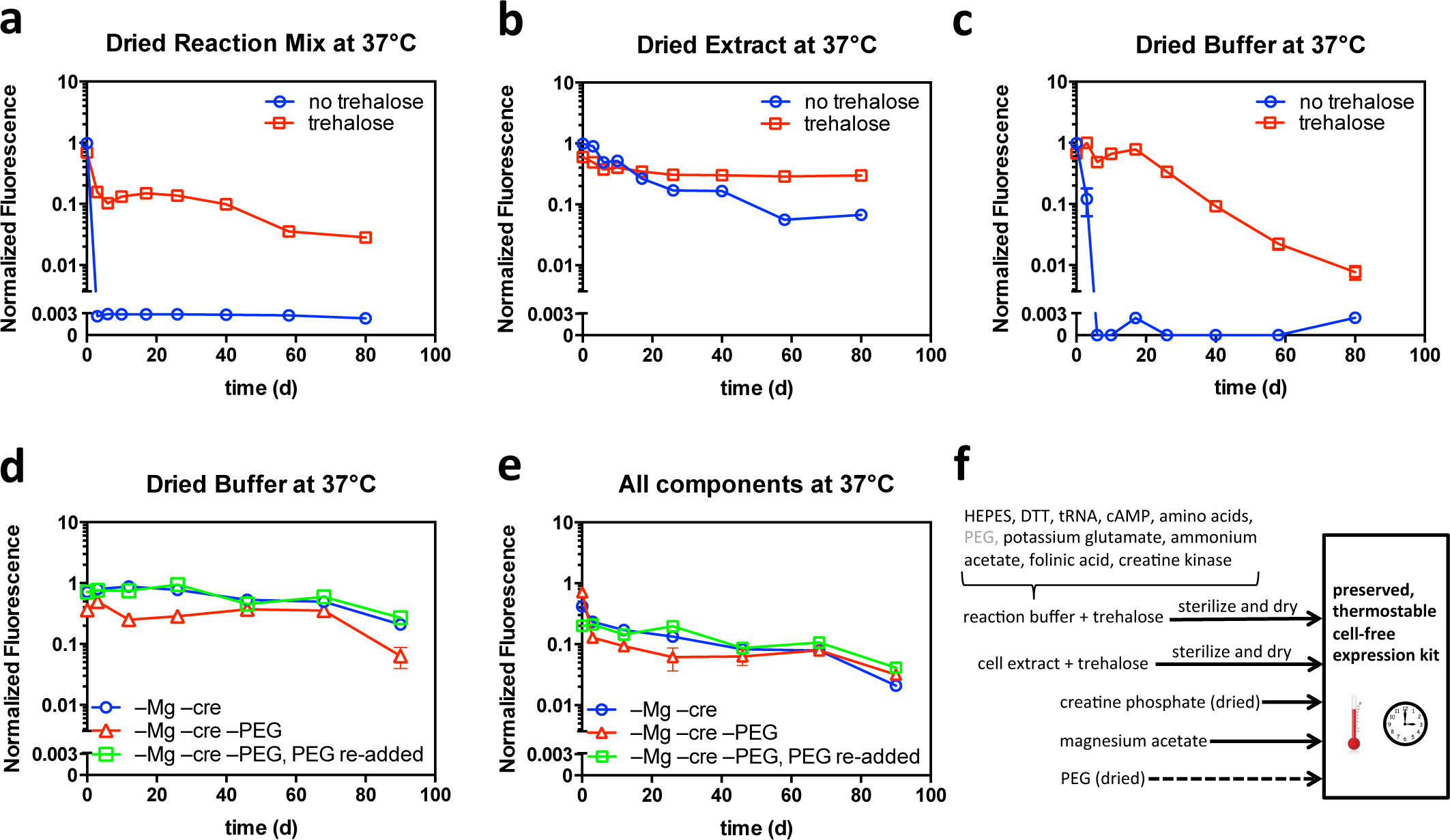
Development of preservation procedure. In all cases, the expression capacity of different reconstituted component combinations was assayed by adding a T7-EGFP expression construct and measuring fluorescence after 5 h of incubation. Fluorescence values are normalized by dividing by the 5 h fluorescence of a standard reaction with fresh reagents. The means of triplicate measurements are shown, with error bars indicating standard deviation. Cases where no error bars are visible indicate that error bars are smaller than the marker. The axis break indicates the threshold for significance above background levels. (**a**) Aliquots of reaction mixture were prepared with and without trehalose and were dried. At different time points during 37°C storage, water was added to reconstitute dried reaction mixture, magnesium acetate was added, and expression capacity was assayed. (**b**) Aliquots of cell extract were prepared with and without trehalose and were dried. At different time points during 37°C storage, water was added to reconstitute dried cell extract, fresh reaction buffer was added, and expression capacity was assayed. (**c**) Aliquots of reaction buffer were prepared with and without trehalose and were dried. At different time points during 37°C storage, water was added to reconstitute dried reaction buffer, fresh cell extract and magnesium acetate were added, and expression capacity was assayed. (**d**) Two variants of reaction buffer were prepared: one omitting creatine phosphate and magnesium acetate and one omitting creatine phosphate, magnesium acetate, and PEG. At different time points during 37°C storage, water, magnesium acetate, and creatine phosphate were added to reconstitute the two different dried reaction buffer variants. A third reaction buffer sample was created by adding PEG to an aliquot of the reconstituted reaction buffer that was dried without PEG. For all samples, fresh cell extract was added, and expression capacity was assayed. (**e**) Cell extract was preserved and tested with the same reaction buffer preparations as in (d). Thus, all components of the cell-free system were preserved and heat stressed under atmospheric conditions. (**f**) Summary of cell-free preservation approach.

While simply drying cell extract improved stability, the significant decreases in the efficacy of extract and reaction buffer over a month suggested the need for identifying preserving agents. For this, we selected the non-reducing sugar alcohol, trehalose. We chose trehalose because it is implicated in anhydrobiosis in natural organisms such as the tardigrade (water bear or moss piglet)^22^, has been widely used in preservation applications^23^, and can be produced in a cost effective manner^24^. As expected, trehalose dramatically extended the shelf life of preserved cell-free systems. Whereas reaction mixture dried without trehalose lacked viability after 1 d, reaction mixture dried with trehalose did not undergo a full 10-fold reduction in yield until after 40 d at 37°C (Fig. 2a). Cell extract efficacy was similarly stabilized over longer time periods (over three months) when dried with trehalose (Fig. 2b), as was dried reaction buffer which, with the addition of trehalose, maintained full stability for 20 d before gradually declining (Fig. 2c).

Although the results in Figure 2b-c demonstrated efficient preservation of cell extract, they showed room for improvement for reaction buffer preservation. Therefore, we explored the drying of different combinations of reaction buffer components. Although we tested several reagents (e.g. PEG as shown in Fig. 2d-e; see also Supplementary Note 1 and Supplementary Fig. 2), only the phosphate donor (here, creatine phosphate) was identified as a reagent that should be preserved separately (Supplementary Figs. 3-4). Separate preservation of the phosphate donor yielded a major improvement in long-term reaction buffer efficacy, particularly after 20 d (Fig. 2d vs. Fig 2c).

Having demonstrated successful expression from preserved cell extract and fresh reaction buffer and from preserved reaction buffer and fresh cell extract, we next tested system performance when preserved cell extract is mixed with preserved reaction buffer. Figure 2e shows expression yield vs. storage time at 37°C for cases in which all expression components (extract, reaction buffer, creatine phosphate, and magnesium acetate) are preserved, heat stressed, and reconstituted (see also Supplementary Fig. 5). In all cases, expression yield is initially lower when preserved extract is used (Fig. 2e) as opposed to fresh extract (Fig. 2d). This is at least partially due to the fact that, when both extract and reaction buffer components are preserved, reconstituted, and combined, the final concentration of trehalose is higher – a consequence of including trehalose in both the extract and the buffer preservation procedures. High concentrations of trehalose can lower protein expression rates (Supplementary Fig. 6). Nonetheless, significant expression is realized for over three months of storage. In a similar experiment for a longer time period, we found that, although expression begins to decrease after approximately two months of exposure at 37°C, significant fluorescence resulting from EGFP expression is realized over eight months of exposure at 37°C (Supplementary Fig. 7). More specifically, full efficacy is still observed at 38 d at 37°C. After this point, expression yield decreases and eventually plateaus to approximately 10%. This degree of heat stability is unprecedented.

Figure 2f summarizes our overall cell-free preservation approach: 1) cell extract with added trehalose is sterilized and dried, 2) reaction buffer prepared without creatine phosphate, without magnesium acetate, with trehalose, and optionally with PEG is sterilized and dried, 3) creatine phosphate and magnesium acetate are then stored separately, 4) all reagents are reconstituted with water, mixed in the appropriate ratios, and then combined with a chosen DNA construct to express the protein of interest. As an alternative to this preservation protocol, Figure 2a shows that the simple drying of reaction mixture combined with trehalose along with the separate storage of magnesium acetate also constitutes a viable approach to preservation. Nonetheless, the approach in Figure 2f offers flexibility advantages. Namely, different buffer variants could be selected at the time of reconstitution to optimize expression for particular protein(s).

Having established a cell-free reagent preservation approach, we sought to demonstrate therapeutic application potential. For this, we used our preserved protein expression system to produce pyocin S5, a 56.1 kDa protein, which, upon proper folding, forms lethal pores in susceptible strains of the pathogen *P. aeruginosa*^25, 26^. To test the long-term therapeutic efficacy of our system, we used both plate clearing and broth dilution assays. The plate clearing experiments in Figure 3a,c show that complete clearing was achieved at all time points throughout the 136 d of heat stress. Likewise, the broth dilution experiments (Figure 3b,d) show that the survival fraction of *P. aeruginosa* was reduced by 4 logs or more (depending on trehalose concentration) using a 20-fold dilution of pyocin produced from 136-day-old, heat-stressed (37°C) cell-free reagents (extract, reaction buffer, magnesium acetate, and creatine phosphate). Impressively, throughout the 136 d experiment a 100-fold dilution of pyocin produced from reconstituted cell-free reagents was sufficient to achieve a 100-fold reduction in the *P. aeruginosa* survival fraction. To further quantify these results, we calculated MIC values and found that they ranged within an order of magnitude for the duration of the 136-day experiment (Supplemental Note 2, Supplementary Figure 8).

**Figure 3:**
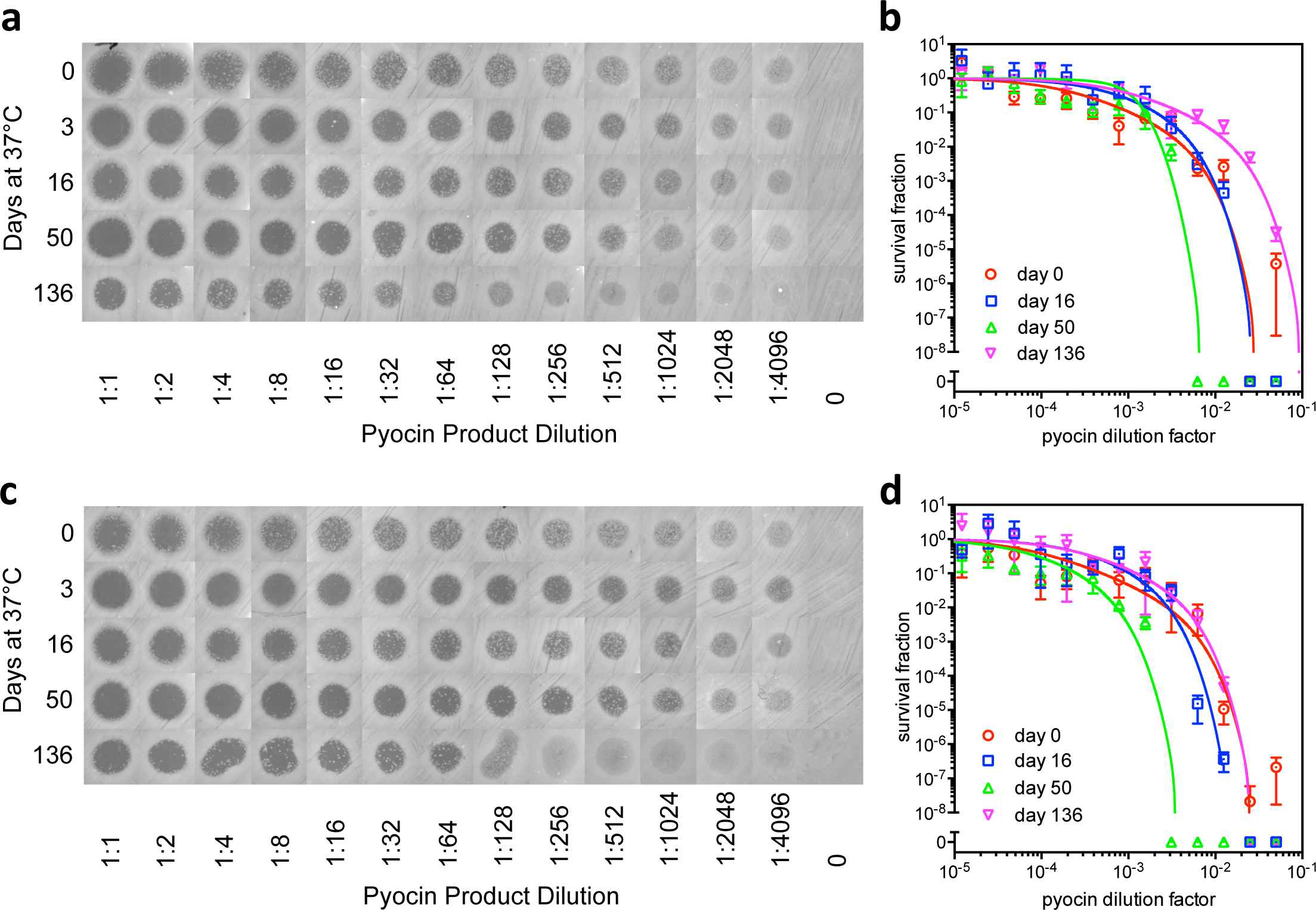
Production of active, therapeutic protein using preserved, heat stressed cell-free protein synthesis reagents stored under atmospheric conditions. Reagents were prepared according to Figure 2f, with PEG omitted from the reaction buffer before drying and not added following reconstitution. At different time points during 37°C storage, reagents were reconstituted and combined. Pyocin S5 and EGFP were expressed in separate reactions. Two experiments were conducted using variants of dried reaction buffer prepared with slightly different trehalose concentrations. (**a**) Clearing assay for visualizing killing efficiency of *P. aeruginosa* and (**b**) broth dilution assay (see Supplementary Note 2) for quantifying killing efficiency. In (a,b), we used reaction buffer prepared to 2.7 times the final reaction concentration with 0.53 M trehalose prior to drying. (**c**) Clearing assay for visualizing killing efficiency of *P. aeruginosa* and (**d**) broth dilution assay (see Supplementary Note 2) for quantifying killing efficiency. In (c,d) we used reaction buffer prepared to 2 times the final reaction concentration with 0.6 M trehalose prior to drying. In (b,d), data points indicate the mean of triplicate plate counts, error bars indicate standard deviation, and curve fits are described in Supplementary Note 2.

## DISCUSSION

Taken together, our results demonstrate a significant advancement in preservation of protein expression systems for field applications. Our approach consists of separately storing cell extract dried with trehalose, dried energy source (creatine phosphate), magnesium acetate, and remaining reaction buffer components dried with trehalose. Preserved components exhibit unprecedented shelf life under heat stress and atmospheric conditions, as we achieved detectable production of EGFP and sufficient pyocin to kill the pathogen *P. aeruginosa*, even after 136 d of exposure of all essential cell-free reagents to 37°C heat stress.

For our demonstrations, we chose a simple, cell-free extract production protocol, and we developed an open-air drying method, making our procedures possible without the need for expensive lyophilization equipment. Future work of extending our methods of trehalose usage and strategic separation of components to drying approaches like lyophilization will be important for large-scale operations. An additional potential improvement of our approach includes the use of antibiotics for reagent sterilization. Here, we chose membrane filtration simply to avoid potential interference with our application demonstration of killing bacteria. However, sterile filtration has been previously associated with long term decreases in cell-free expression yield^27^. Therefore, alternative sterilization approaches may not only lend themselves towards better scalability^28^, but also further improve preservation^28^.

Our demonstrated, long-term heat stability of protein expression opens the door to a host of applications in the realms of therapeutics production and delivery, bioremediation, and biosensing. Of particular relevance, the potential end of the antibiotic era has brought increased attention to the production of alternative therapeutics^16^, including pyocin‐ and colicin-like proteins and more complicated targets like whole phages^29^, antibiotic peptides^30^, and vaccine components^31^. Previous studies have demonstrated efficient cell-free production of these types of therapeutics, and recently, novel fluidics platforms have been developed to facilitate the expression and purification of proteins using cell-free systems^32, 33^. Our work fills a key need to make practical use of these cell-free capabilities and platforms, as it addresses the limiting factors of stability, storage, and distribution associated with protein therapeutics, and enables production of these novel antimicrobials on site.

## METHODS

### Cell extract

To prepare cell-free expression reagents for demonstration experiments, a method similar to that of Kim et al was used^20^. Specifically, *E. coli* BL21 Star (Invitrogen) cells were cultured overnight in 100 mL 2xYPTG media in a 500 mL flask at 37°C in a shaking incubator (225 RPM). 5 mL of this starter culture was used to inoculate 500 mL 2xYPTG in 2 L flasks at 37°C and 225 RPM shaking. In order to induce expression of T7 RNA polymerase, cultures were induced with 1 mM IPTG when the OD_600_ reached approximately 0.6. Meanwhile, 3 L of ‘Buffer A’ was prepared in RNAse cleaned glassware and was chilled on ice. Buffer A consisted of 10 mM Tris–acetate buffer (pH 8.2), 14 mM magnesium acetate, 60 mM potassium glutamate, 1 mM dithiothreitol (DTT), and 0.05% 2-mercaptoethanol.

Cells were harvested in mid-log phase by centrifugation at 4000*g* for 20 min at 4°C. Wet cell pellet masses were measured. The cells were then washed three times by suspension in 20 mL of Buffer A per gram of wet cells and subsequent centrifugation at 4000*g* for 10 min. Following the three washes, cell pellets were weighed again and were stored at -80°C overnight. The next day, 100 mL of ‘Buffer B’ was prepared, which consisted of Buffer A without mercaptoethanol, and was chilled on ice. The cell pellets were thawed by placing their containers in room temperature water. Just before the cells were completely thawed, they were transferred to ice to keep the cells below 4°C. The thawed cells were then suspended in 1.27 mL of Buffer B per gram of cell mass. The resuspended cells were then disrupted by sonication for 10 minutes on ice. The sonicated lysate was centrifuged for 10 minutes at 12,000*g*. Supernatants were transferred to fresh centrifuge tubes and again centrifuged for 10 minutes at 12,000*g*. Supernatants were again transferred to fresh centrifuge tubes and were centrifuged for 10 minutes at 25000*g*. The final supernatants were transferred to RNAse free 50 mL conical tubes and were incubated for 30 minutes at 37°C. The extract was then divided into small aliquots and stored at −80°C.

### Reaction buffer

The standard reaction buffer was based on previous protocols ^20^. Reaction buffer was set up as a two-fold concentrated mixture, such that a cell-free expression reaction with 50% reaction buffer would have the following reagents at the specified final concentrations:

28.5 mM Hepes–KOH (pH 8.2)
1.2 mM ATP
0.85 mM each of CTP, GTP and UTP
2 mM DTT
0.17 mg/mL E. coli total tRNA mixture (from strain MRE600)
0.64 mM cAMP
90 mM potassium glutamate
80 mM ammonium acetate
24 mM magnesium acetate
34 μg/mL L-5-formyl-5,6,7,8-tetrahydrofolic acid (folinic acid)
4mM of cysteine and 2.1 mM of each of 19 other amino acids
2% PEG (8000)
67 mM creatine phosphate
3.2 μg/mL creatine kinase (CK)

### Plasmids

The EGFP expression construct pUCT7tet-T7term expresses EGFP from a TetR repressible T7 promoter variant and has been previously described^7^. The pyocin expression construct pT7-pyoS5, expresses pyocin S5 from *P. aeruginosa* PAO-1 from a T7 promoter. More specifically, it contains a T7 promoter, a strong ribosome binding site, the *pyoS5* gene, and a T7 terminator in a modified pUC19 backbone. This plasmid was constructed using DNA synthesized by GeneArt and assembled through Seamless Cloning reactions.

### GFP expression assays

Triplicate 20 μL reactions were set up in the wells of a black, 384-well microplate, and 20 μL of mineral oil was added to prevent evaporation. Immediately following set up, reactions were run in a Tecan Saffire II microplate reader with incubation at 35°C, fluorescence measurements every five minutes, and two minutes of shaking in between measurements. Fluorescence was captured using 483 nm (20 nm bandwidth) excitation, 525 nm emission (20 nm bandwidth), and gain set at 55. Background correction was performed in order to capture fluorescence resulting from EGFP as opposed to the cell-free reagents. To estimate background, robust, locally-weighted polynomial regression with a span of four samples was performed in Matlab to smoothen the data. This smoothening process was aimed at minimizing transient artifacts. The artifacts may arise from a number of potential sources, including instrument noise, oil bubbles moving/settling, and initially incomplete mixing of the viscous cell-free reagents. While these transients are generally small, the background fluorescence is also small, necessitating their suppression. Background for each well measurement was then calculated as the minimum of the first three readings, since the earliest readings capture fluorescence prior to transcription, translation, and folding of EGFP. Expression yield was then estimated as the fluorescence after 5 h of incubation, with the background fluorescence subtracted. Despite efforts to correct for background, instrument noise, artifacts, and a gradual increase in background fluorescence of cell-free reagents contribute to a detection limit, below which fluorescence from EGFP expression cannot be distinguished from background. To estimate this detection limit, we performed the experiments from Figure 1b-d after a week of storage with no DNA added. The maximum background corrected value obtained from this experiment, plus one standard deviation was set as the detection limit. In all plots, we set an axis break to delineate this detection threshold.

### Baseline stability characterization

To establish an initial preservation performance baseline, we conducted experiments to determine the inherent shelf life of cell-free reaction mixture (Fig. 1). To characterize reaction mixture stability, we first sterilized reaction mixture through 0.22 μm filtration to prevent initial fouling that may obscure inherent biochemical stability. Aliquots of filtered reaction mixture were then stored at room temperature and at 37°C. At different time points, the plasmid pUC-T7tet-T7term was added to express enhanced green fluorescent protein (EGFP) from a T7 promoter variant. Using a microplate reader, the fluorescence intensity after 5 h of incubation was measured to quantify expression, as described in the GFP expression assay section above.

To gain a deeper understanding of reaction mixture degradation, we also characterized cell extract and reaction buffer stability separately. To quantify cell extract stability, we stored aliquots at room temperature and at 37°C. At different time points, we added fresh reaction buffer and the pUC-T7tet-T7term expression construct and assayed fluorescence as was done to test the full reaction mixture (Fig. 1c). After 4 d at room temperature, cell extract efficacy had decreased by a factor of 31.1. After only one day under heat stress, the extract completely lost viability and generated negligible fluorescence in our expression assay. To test reaction buffer stability, we similarly added fresh extract and the expression construct and quantified fluorescence at different time points (Fig. 1d). At room temperature, buffer efficacy remained high for 43 d and then rapidly decreased. However, after only 9 d at 37°C, the buffer efficacy was reduced by a 41.5-fold.

### Preservation and reconstitution

We utilized a simple open-air drying approach. This approach was chosen due to its simplicity and low cost. Also, the fact that the final product is not trapped in a porous material such as paper facilitates transport and large-scale applications. All of the following approaches could be adapted to other drying methods, such as freeze-drying or spray-drying, if desired. Prior to drying, all reagents were filtered using 0.22 μm spin filters (Millipore). All drying was performed by pipetting small aliquots (14-28 μL) on a silicon sheet (Silpat). The sheet was then placed in a 37°C incubator. Dried samples were recovered using a razor and transferred to DNAse/RNAse free 1.5 mL microcentrifuge tubes. Temperature and humidity were quantified using an Inkbird THC-4 digital sensor. For instance, for the experiments in Figure 3, the average temperature was 37.1°C (σ=0.8°C), and the average relative humidity was 20.5% (σ=4.3%).

Trehalose (Sigma) concentrations between 0.5 M and 0.6 M were chosen to balance preservation efficiency with expression efficiency, since high concentrations of trehalose can inhibit protein expression (Supplementary Fig. 6). Samples in experiments in Figure 2a-c were dried with 0.55 M trehalose as indicated. For the experiments in Figure 2d-e, reaction buffer samples were dried with 0.59 M trehalose, and cell extract was dried with 0.55 M trehalose. Creatine phosphate (Roche) and PEG (Fisher) were stored at 37°C in powder form (as received from manufacturer) and were added to the indicated samples upon reconstitution. For the experiments shown in Figures 2c, 2d, 2e, and 3, a 1 M solution of magnesium acetate was prepared, filter sterilized, and stored at 37°C for the duration of each experiment. While dry powder could have been used, this approach facilitated small scale testing.

### Pyocin killing

Reaction buffer without PEG was preserved, and PEG was not added upon reconstitution. Two similar experiments were conducted simultaneously, with variants of reaction buffer prepared with different trehalose concentrations. Specifically, for the experiment in Figure 3a-b, the initial reaction buffer was prepared to 2.7 times the final reaction concentration with 0.53 M trehalose prior to drying. For the experiment in Figure 3c-d, which was performed at the same time, the initial reaction buffer was prepared at 2 times the final reaction concentration with 0.6 M trehalose prior to drying. In all cases, cell extract (dried with trehalose), magnesium acetate, and creatine phosphate were heat stressed at 37°C and stored under atmospheric conditions along with the reaction buffers. At different time points, we reconstituted and combined all expression components to perform protein expression reactions. Pyocin S5 was expressed using the T7 expression plasmid pT7-pyoS5, and EGFP was expressed using pUCT7tet-T7term. These reactions were incubated in microcentrifuge tubes in a rotator at 30°C. After 6 h, different dilutions of the pyocin reaction were prepared, whereby the EGFP reaction product was used to dilute the pyocin reaction product.

For each time point in Figure 3, 14 samples were set up for analysis by plate clearing and broth dilution assays. Of these 14 samples, 13 consisted of two-fold serial dilutions of the pyocin cell-free reaction product, using the cell-free EGFP reaction product to dilute the pyocin product. The final sample consisted of the cell-free EGFP reaction product, which served as a negative control. Broth microdilution methods were adapted from Wiegand et al^34^, and plate clearing assays were based on a combination of the Kirby Bauer method and previously used assays for characterizing pyocin S5^25^. For both plate clearing and broth dilution, three colonies of *P. aeruginosa* BE171^35^ were cultured in 2 mL casamino acids (CAA) media (5g/L Difco iron-poor casamino acids, 0.25 g/L MgSO_4_*7H_2_O, and 1.18 g/L K_2_HPO_4_) and grown to log phase in a 37°C shaking incubator. Cultures were diluted in CAA to prepare a 2 mL stock with an OD600 of 0.1. For the plate clearing assays, petri dishes (100 mm diameter) were prepared with 25 mL CAA and 1.5% agar. These CAA plates were inoculated by swabbing approximately 200 μL of culture diluted to an OD_600_ of 0.1 with a sterile cotton tipped applicator. Once the swabbed solution dried, 10 μL drops of each of the 14 cell-free reaction samples were dispensed (7 samples per plate). Plates were then incubated at 37°C for approximately 15 h and subsequently imaged using a G:Box Chemi XX9 imager. For broth dilution assays, a bacterial suspension of 5*10^5^ CFU/mL was prepared, and 95 μL of this suspension was aliquoted to wells of a 96 well microplate. Then, 5 μL of each cell-free reaction sample was added to the cell suspension in wells. An additional control was set up, consisting of 95 μL of sterile media and 5 μL of the thirteenth serial dilution. The microplate was then incubated at 37°C with shaking for approximately 16 h, and survival (CFU/mL) was then quantified using the Miles and Misra method.^36^

## ACKNOWLEDGEMENTS

This research was supported by the Independent Research and Development Program of the Johns Hopkins University Applied Physics Laboratory. Preparation of the manuscript was funded by the Johns Hopkins Applied Physics Laboratory Janney Publication Program (to D.K.K.). We thank E.P. Greenberg for providing the BE171 strain of *P. aeruginosa* (USPS P30DK089507), M.J. Doktycz and J.L Morrell-Falvey for sending useful materials (pUCT7tet-T7term), Z. Haque for assistance during early stages of the project, and J. Hamel, J.D. Evans, and P.A. Demirev for helpful feedback and discussions.

## AUTHOR CONTRIBUTIONS

SZ, SB, and DKK performed experiments. DKK analyzed data, designed experiments, and wrote the manuscript. PT, JW, and DKK discussed results and edited the manuscript.

